# Predicting skeletal stature using ancient DNA

**DOI:** 10.1101/2021.03.31.437877

**Authors:** Samantha L Cox, Hannah Moots, Jay T Stock, Andrej Shbat, Bárbara D Bitarello, Wolfgang Haak, Eva Rosenstock, Christopher B Ruff, Iain Mathieson

**Affiliations:** Department of Genetics, Perelman School of Medicine, University of Pennsylvania, Philadelphia, PA; Physical Anthropology Section, Penn Museum, University of Pennsylvania, Philadelphia, PA; Department of Anthropology, Stanford University, Stanford, CA; Department of Anthropology, Western University, London, Ontario, Canada; Department of Archaeology, Max Planck Institute for the Science of Human History, Jena, Germany; Institute of Anatomy, First Faculty of Medicine, Charles University, Prague, Czech Republic; Department of Archaeogenetics, Max Planck Institute for the Science of Human History, Jena, Germany; Einstein Center Chronoi, Freie Universität Berlin, Berlin, Germany; Center for Functional Anatomy and Evolution, Johns Hopkins University School of Medicine, Baltimore, MD

## Abstract

**Objectives:** Ancient DNA provides an opportunity to separate the genetic and environmental bases of complex traits by allowing direct estimation of genetic values in ancient individuals. Here, we test whether genetic scores for height in ancient individuals are predictive of their actual height, as inferred from skeletal remains. We estimate the contributions of genetic and environmental variables to observed phenotypic variation as a first step towards quantifying individual sources of morphological variation.

**Materials and Methods:** We collected stature estimates and femur lengths from West Eurasian skeletal remains with published genome-wide ancient DNA data (n=167, dating from 33,000-850 BP). We also recorded genetic sex, genetic ancestry, date and paleoclimate data for each individual, and *δ*^13^C and *δ*^15^N stable isotope values where available (n=67).

**Results:** A polygenic score (PRS) for height predicts 6.8% of the variance in femur length in our data (n=117, SD=0.0068%, p<0.001), controlling for sex, ancestry, and date. This is consistent with the predictive power of height PRS in present-day populations and the low coverage of ancient samples. Comparatively, sex explains about 15% of the variance in femur length in our sample. Environmental effects also likely play a role in variation, independent of genetics, though with considerable uncertainty (longitude: *R*^2^=0.0317, SD=0.009, p=0.019).

**Discussion:** Polygenic scores explain a small but significant proportion of the variance in height in ancient individuals, though not enough to make useful predictions of individual phenotypes. However, environmental variables also contribute to phenotypic outcomes and understanding their interaction with direct genetic predictions will provide a framework with which to model how plasticity and genetic changes ultimately combine to drive adaptation and evolution.

## Introduction

One of the central goals of biological anthropology is to understand the environmental and genetic contributions to phenotypic change. Skeletal series covering long time periods and diverse environments are informative, but limited by inability to separate these contributions—a limitation that can now be addressed with ancient DNA. The ability to generate genetic data from skeletal remains has had an enormous impact on studies of human history. By identifying genetic links among individuals and populations, ancient DNA allows us to reconstruct demographic histories on both large and small scales (Skoglund & Mathieson, 2018; Racimo et al., 2020), as well as the effects of natural selection (Marciniak & Perry, 2017). To learn about complex trait evolution, ancient DNA can be combined with information from genome-wide association studies (GWAS). These studies, now involving millions of individuals, have identified large numbers of single-nucleotide polymorphisms (SNPs) associated with hundreds of phenotypes (Visscher et al., 2017). Though the effect of any individual variant is typically small, the effects of many SNPs can be summed to produce a polygenic score (PRS) which can be thought of as an estimate of an individual’s genetic potential or risk for a particular phenotype (Torkamani et al., 2018).

Height is a polygenic trait which is highly heritable and easy to measure. GWAS have identified thousands of SNPs significantly associated with height (Yengo et al., 2018). Each one has only a tiny effect—±1-2mm per SNP—but combined explain around 25% of the phentoypic variance in present-day populations of European ancestry (Yengo et al., 2018). Contingent upon reasonable preservation of skeletal remains, stature estimation from long bones is relatively straightforward, and there is an excellent record of changes in stature in many parts of the world (Bogin & Keep, 1999; Cohen & Crane-Kramer, 2007; Ruff, 2018; Rosenstock et al., 2019). On a population level, changes in polygenic score in Europe computed from ancient DNA largely track trends in stature in the European skeletal record (Cox et al., 2019). However, environmental effects on stature can still be large, as shown by 20*^th^* century secular trends (NCD Risk Factor Collaboration, 2016), and are not confined to recent history. For example, during the Bronze Age, genetic predictions suggest increasing stature, but estimated skeletal heights actually decrease (Cox et al., 2019).

Polygenic scores have three main limitations. First, due to incomplete correction of population stratification in GWAS, they can capture environmental variation in present-day populations leading to ancestry-related biases (Sohail et al., 2019; Berg et al., 2019). Second, their accuracy decreases as genetic distance from the present-day European ancestry GWAS populations increases (Martin et al., 2019). Finally, their accuracy can be reduced by environmental differences, even within-population (Mostafavi et al., 2020). We therefore set out to test whether polygenic scores based on present-day GWAS predict height in ancient individuals by collecting data for individuals with both ancient DNA and skeletal measurements. This allows us to assess the validity of complex trait prediction in ancient individuals, and whether we can use this approach to understand the relationship between genetic and environmental components of stature and, perhaps, of other phenotypes.

## Materials and Methods

### Data Collection

We collected genetic data from published ancient DNA (aDNA) studies (31,000-850BP). Most individuals had pseudo-haploid genotypes at a set of 1.24 million SNPs (the “1240k array”) and for individuals with shotgun sequence data we randomly selected a single read at each of the covered sites. The majority of the available ancient DNA data comes from Western Eurasia and so we focused on this region, broadly defined (including individuals from up to 100°E longitude).

For each individual with published aDNA, we attempted to find data on their skeletal measurements. Some aDNA papers include stature and/or femur length measurements in their supplemental materials. For other individuals, we searched archaeological and anthropological literature for published data. However, the vast majority of published aDNA data come from skeletal individuals which are either unpublished or highly fragmented and therefore unmeasurable. We also report new measurements for 30 individuals (Supplementary Table 1).

For each individual, we recorded maximum femur length, when available, otherwise we recorded the estimated stature and estimation method. For individuals with published femur lengths, we estimated stature using the method of Ruff et al. (2012). We restricted analyses to adult individuals free from reported major pathology. Ultimately, from approximately 4000 published aDNA samples (Martiniano et al., 2016; Mittnik et al., 2019; Mathieson et al., 2018; Olalde et al., 2019; de Barros Damgaard et al., 2018; Mittnik et al., 2018; Fu et al., 2016; Schiffels et al., 2016; Narasimhan et al., 2019; Lipson et al., 2017; Olalde et al., 2018; Brace et al., 2019; González-Fortes et al., 2017; Furtwängler et al., 2020; Fernandes et al., 2018; Antonio et al., 2019; Sikora et al., 2017; Krzewińska et al., 2018; Margaryan et al., 2020; Mathieson et al., 2015; Schroeder et al., 2019), we compiled metric data for 167 individuals (Caffell & Holst, 2012; Tebelškis & Jankauskas, 2002; Alpaslan-Roodenberg, 2001; Dunwell, 2007; Andrews & Thompson, 2016; Köhler et al., 2017; Fokkens et al., 2017; Boroneant, 2010; Pardini, 1977; Frei et al., 2019; Kjellström, 2005; Cairns, 2015; Price et al., 2016; Szczepanek, 2013; Alciati, 1967; Massy, 2018; Malmström et al., 2019; Berthold et al., 2008; Kitti, 2008; Saag et al., 2020; Auerbach & Ruff, 2004; Auerbach, 2004; Schiffels et al., 2016; Rosenstock et al., 2019). We removed 28 samples with more than 95% missing genetic data and one individual with an unusually short femur (Fig. 1B), bringing the sample size to 138. Finally, 21 stature estimates used unknown methods and we removed them from the majority of tests, bringing the final sample size for most analyses to 117 (Supplementary Table 1).

**Figure 1:**
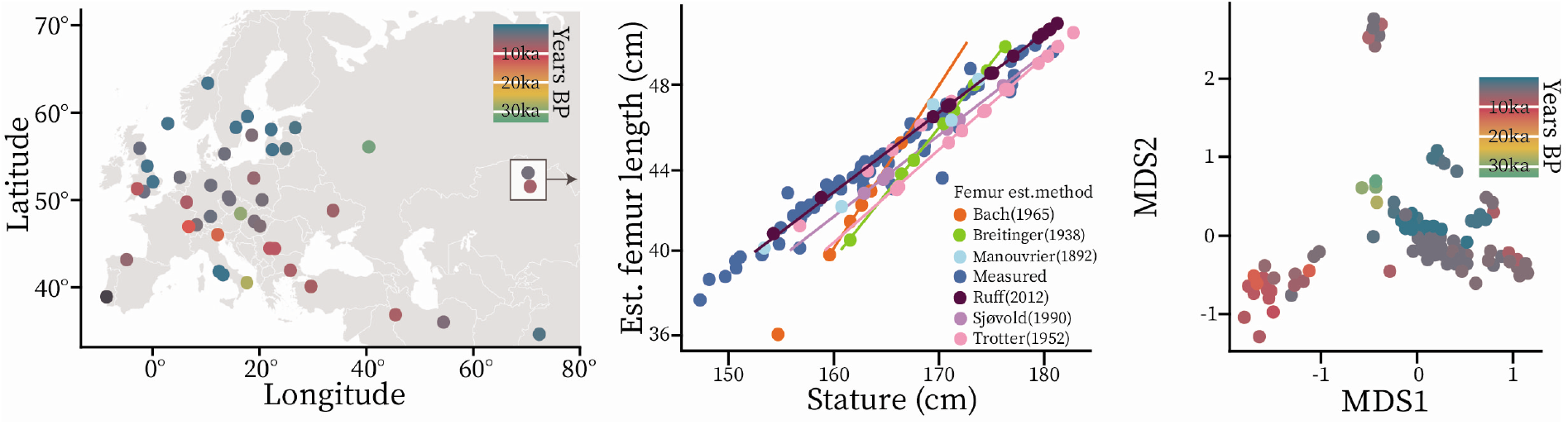
A) map of sites with specimens with both ancient DNA and metric data; B) The relationship between skeletal statures and femur length. Lines indicate the regression line for each stature estimation method. The stature for the outlying individual in the lower left was estimated using the Bach (1965) method, and was removed from further analysis. For all individuals with directly measured femora, stature was estimated using the method by Ruff et al. (2012); C) The first two MDS axes of the genetic data summarizing the genetic ancestry of the samples. Clusters correlate with time, and are largely associated with substantial shifts in genetic ancestry. The cluster in the top right corner represents the most eastern individuals in our sample, from present-day Siberia.

There are a number of methods available for estimating living statures based on skeletal measurements. Population-specific methods are the most accurate, but are not available for every population (Ruff et al., 2012). Researchers ideally choose a method which has been developed on a population similar to that under investigation in terms of ancestry or body proportions; however, this is often not possible and there are a few methods which are most frequently used, even if not population-specific (e.g Trotter & Gleser, 1952; Sjøvold, 1990; Ruff et al., 2012). Ideally we would compare statures estimated using the same equations but that was not possible since in many cases underlying data were not available.

We dealt with this issue in two ways. First, we included stature estimation method as a discrete factor in the linear regression to predict stature, but we were concerned about the proportion of variance attributable to the estimation method in our models and the lack of population-specific equations. Due to this, we took a second approach: since the single bone which offers the most accurate stature estimate is the femur (Trotter & Gleser, 1952; Ruff et al., 2012; Ruff, 2018) we tried predicting maximum femur length rather than stature. Femur lengths were only published for 78 out of our 167 individuals. For the other individuals, since statures are estimated as linear functions of long bone lengths, we inverted the estimation equations to retrieve femur measurements corresponding to each individual stature (Table 1; for further discussion on this approach see Köpke & Baten (2005); Rosenstock et al. (2019)). For individuals for which stature had been estimated using a non-femur long bone, this procedure gives us the femur length which would have produced the originally estimated stature (Fig. 1B). We confirmed that estimation method did not have a significant effect on femur lengths estimated using this approach (P=0.539). We used femur length rather than estimated stature for most analyses, omitting (n=21) individuals for which the stature estimation method was not cited. Finally, the slope of the regression line from which the Bach (1965) formula is derived deviates from those of the other estimation methods and produces outlier femur measurements (Fig. 1B). The applicability of the Bach (1965) method has also been questioned by other researchers (Sládek et al., 2015). This method was only used for a few individuals (n=3), and as inclusion of these individuals did not affect prediction results, all were included in the analysis except for one. This individual (DNA sample ID: AITI_95, a genetic female from the Bronze Age German site of Kleinaitingen-Gewerbegebiet Nord) was estimated to have had an unusually short femur (approx 36cm, estimated stature = 154cm) and was excluded as we could not be confident in the accuracy of the estimate.

**Table 1:** Inverted stature estimation equations

| Reference |  | Stature Equation | Error (cm) | Femur Equation |
| --- | --- | --- | --- | --- |
| Manouvrier (1892)* | Females | $49.319 + 2.543Femur$ | | $\frac{Stature - 49.319}{2.543}$ |
| | Males | $71.065 + 2.13Femur$ | | $\frac{Stature - 71.065}{2.13}$ |
| Pearson (1899) | Females | $72.844 + 1.945Femur$ | | $\frac{Stature - 81.306}{1.88}$ |
| | Males | $81.306 + 1.88Femur$ | | $\frac{Stature - 81.306}{1.88}$ |
| Breitinger (1937) | Males | $94.31 + 1.64Femur$ | $\pm 4.8$ | $\frac{Stature - 94.31}{1.64}$ |
| Bach (1965) | Females | $106.69 + 1.313Femur$ | $\pm 4.1$ | $\frac{Stature - 106.69}{1.313}$ |
| Trotter & Gleser (1952) | White females | $54.10 + 2.47Femur$ | $\pm 3.72$ | $\frac{Stature - 54.10}{2.47}$ |
| | White males | $61.41 + 2.38Femur$ | $\pm 3.27$ | $\frac{Stature - 61.41}{2.38}$ |
| Sjøvold (1990) | Independent | $49.96 + 2.63Femur$ | $\pm 4.52$ | $\frac{Stature - 49.96}{2.63}$ |
| Ruff et al. (2012) | Females | $43.56 + 2.69Femur$ | $\pm 2.92$ | $\frac{Stature - 43.56}{2.69}$ |
| | Males | $42.85 + 2.72Femur$ | $\pm 3.21$ | $\frac{Stature - 42.85}{2.72}$ |

\*These equations are interpolated from the data given in the tables of the original publication.

We also collected other variables for inclusion in our models: ancestry, date, sex, climate, and diet. We estimated genetic ancestry by multi-dimensional scaling (MDS, with k=4 dimensions) of the genetic data—referred to collectively as the “ancestry” variable. We determined the date of each sample based on the calibrated radiocarbon dates reported in the original publication of the genetic data. For the few samples for which there was no direct date available, we used the mid-point of the archaeological date range. Genetic sex was reported in the original publications. Climate variables of mean daily temperature and annual precipitation were obtained from the 5min (medium resolution, data points every 10km) paleoclimate dataset available at PaleoClim.org (Brown et al., 2018). We extracted the relevant data using the *raster* package (Hijmans & van Etten, 2012) in R. Using geographic coordinates, we calculated the distance from the sites where skeletons were excavated to the surrounding climate data points (using the gdist() function from the *Imap* package; Wallace (2012)), and chose the point closest to the site to represent its climate. For Western Europe, most sites are within a few kilometers of available climate data; however, there are a handful of sites in present-day Russia and the Middle East which are quite far from any available PaleoClim data (200-1500km). To represent diet, we searched for stable isotope data (*δ*^13^*C* and *δ*^15^*N*). However, these data were only published for about half the individuals (n=66)(Price et al., 2016; Antanaitis-Jacobs et al., 2009; Stockhammer et al., 2015; Scheibner, 2016; Antanaitis-Jacobs & Ogrinc, 2000; Lehuray et al., 2006; Afshar et al., 2019; Waterman et al., 2016; Weber et al., 2011, 2016; Müldner et al., 2011; Antonio et al., 2019; Szczepanek, 2013; Olalde et al., 2018; Andrews & Thompson, 2016; Dunwell, 2007; Mathieson et al., 2018; Berthold et al., 2008; Malmström et al., 2019; Kjellström, 2005; Kjellström et al., 2009; Scheibner, 2016).

### Polygenic Scores

We calculated PRS using UK Biobank standing height summary statistics calculated by the Neale Lab (2018). After intersecting these sites with those on the 1240k array, we tested a variety of PRS constructions. For the main analysis, we further restricted to HapMap3 SNPs (leaving 405,502 remaining) and estimated SNP weights using the infinitesimal model of *LDpred* (Vilhjalmsson et al., 2015) with an LD reference panel made up of 8,000 randomly chosen individuals of British ancestry from the UK Biobank. We then computed polygenic scores using the --score command in *plink* (Chang et al., 2015). As an alternative approach, we also calculated PRS using a simpler clumping/thresholding approach. Clumping parameters included all combinations of: 10^−2^, 10^−6^, and 10^−8^ p-value cut-offs; 100, 250, and 500 kb windows; and *r*^2^ cutoffs of 0.1, 0.3, and 0.5. We used *plink 1.9* (Chang et al., 2015) to clump (--clump) SNPs using these parameters with an LD reference panel made up of 500 individuals from five European populations (1000 Genomes Project Consortium, 2015), and to compute polygenic scores (--score). In both approaches, missing genotypes are replaced by the sample mean frequency—a conservative approach that shrinks scores towards the sample mean.

### Genotype Imputation

The genetic data are relatively low coverage (haploid median=0.607, range=0.001-1). Therefore it is not possible to infer diploid genotypes, and we use pseudo-haploid data that represents a single allele at each site. This limits performance of the PRS because effectively we only see at most half of each individual’s genotype. In practice samples often perform worse because many sites are missing data entirely. We attempted to improve individual prediction by using genotype imputation to infer diploid genotypes and missing sites. We downloaded bam files for each of the individuals in our sample, and extracted reference/alternative read counts at each of the 1240k sites. We computed genotype likelihoods based on a binomial distribution of reads with a 2% rate of error. We then ran *Beagle4* (Browning & Browning, 2016) using a reference panel made up of the European ancestry populations from the 1000 Genomes Project Consortium (2015) to impute diploid genotypes at all 1240k sites, including those that were missing in the original data. Finally, we set any sites with a genotype imputation accuracy *R*^2^ < 0.8 to missing.

### Statistical Analysis

We fit linear models of femur length (and stature) as a function of sex, PRS, genome-wide ancestry, and date, and also included stable isotope and climate variables when appropriate. The ancestry component includes 4 MDS axes. Date includes both date (years before present) and date squared, to allow for nonlinear effects (Fig. 2B). We evaluated the contribution of each term based on the difference in *R*^2^ between the full model and a reduced model without the term being tested; we refer to this difference simply as *R*^2^. To test the significance of a particular variable, we permuted its value within the dataset, keeping constant those variables which were not being tested. We permuted each test variable 10,000 times. We computed P-values as the proportion of times that the *R*^2^ value in the original data was greater than the *R*^2^ of the permuted distribution. When permuting terms with multiple components, for example ancestry, we ensured that the relationship between each of the permuted components was maintained.

**Figure 2:**
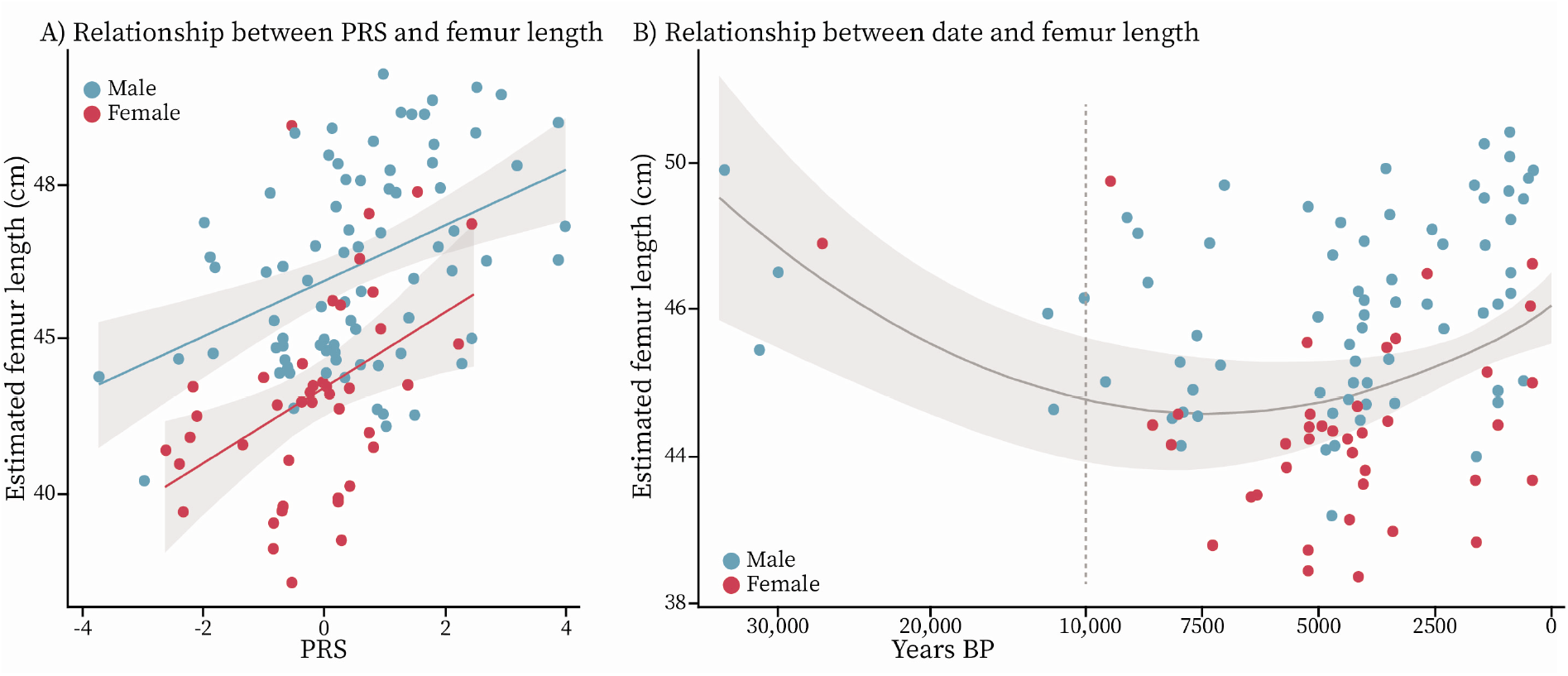
A) Plot of the linear relationship between PRS and femur length. Higher PRS values are associated with longer femur lengths in the data. Colors indicate sex, the lines are the regression lines for males and females separately, and the grey shadows are the 95% confidence intervals. For our main results, we assume the slope of this regression is identical between sexes. B) Plot of the fitted quadratic relationship between date and femur length. Colors indicate sex, the solid grey line is the quadratic fit line for the pooled-sex group, the grey shadow is the 95% confidence interval, and the vertical dashed line indicates the change in x-axis plotting scale.

## Results

The *LDpred* polygenic score predicts 6.8% of the variation in femur length in our data. We fitted linear models including sex, PRS (Fig. 2A), ancestry (4 MDS components) and date (including date squared; Fig. 2B) and evaluated significance by permuting the terms being tested. The *R*^2^ of PRS was 0.068 (SD=0.007, p=0.000) (Table 2), showing that it explains a small but statistically significant proportion of the variance in femur length in our data, once the other variables are taken into account. For comparison, this is less than half of the variance explained by sex (*R*^2^= 0.15). Ancestry had *R*^2^=0.056 (SD=0.0131, p=0.015); a contribution that may also partially reflect genetic effects. In a model without ancestry, the *R*^2^ of the PRS term increases to 0.088, reflecting systematic variation in PRS with ancestry. Date has *R*^2^=0.043 although this largely reflects differences between the Early Upper Paleolithic and later populations (Fig. 2B). When we predicted stature, rather than femur length, including a constant term for estimation method, we found consistent results, though with a slightly lower *R*^2^, possibly due to the error introduced by the variation in estimation methods (PRS *R*^2^=0.048, SD=0.0049, p=0.000) (Table 2).

**Table 2:**
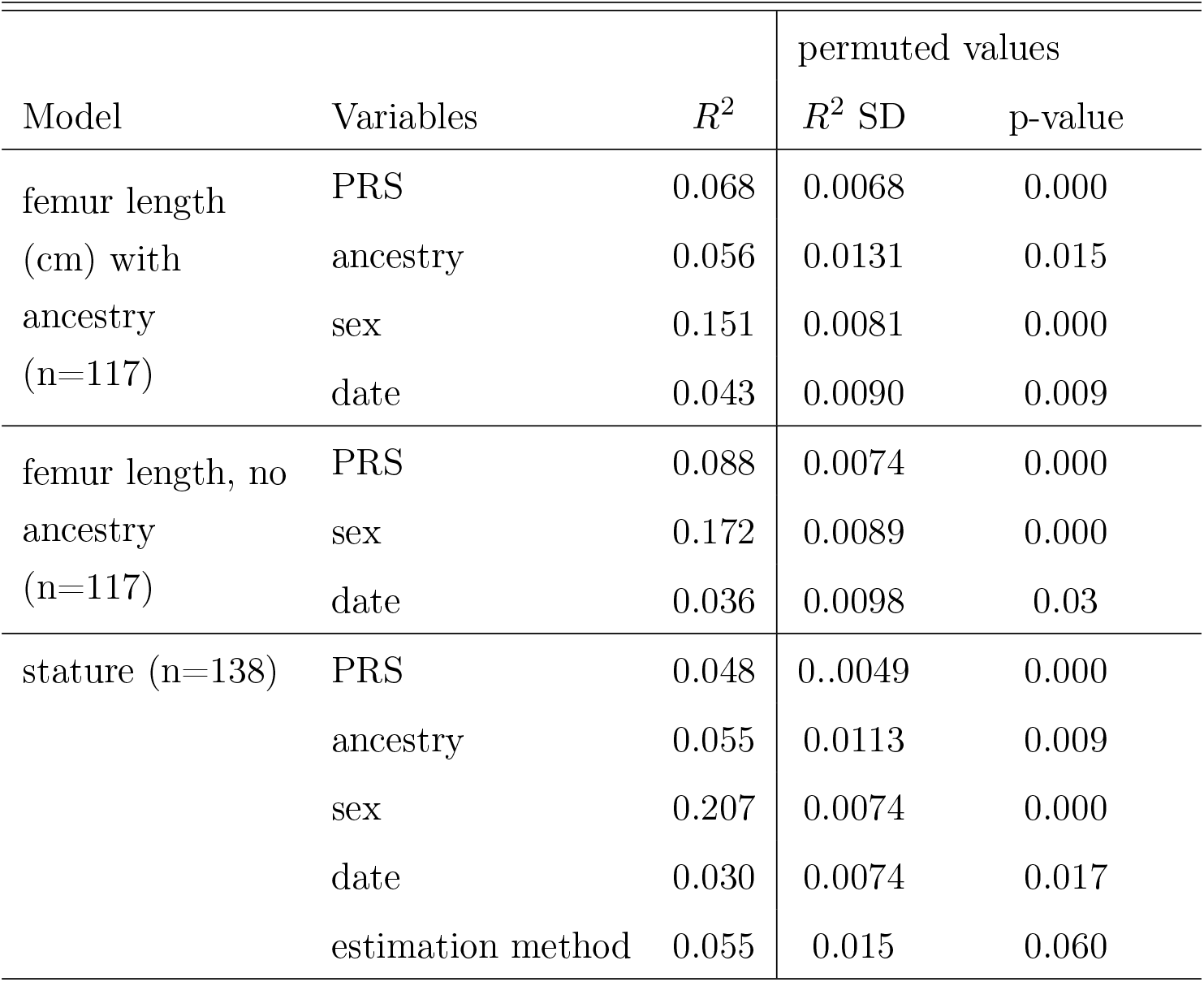
Linear model results

Almost all the predictive power of the PRS comes from samples above median coverage (haploid median coverage=0.63, Fig. 3B&C). To test how PRS construction might affect prediction results, we also constructed PRS using a more traditional clumping and thresholding method. For this, we computed PRS using p-value cut-offs of 10^−2^, 10^−6^, and 10^−8^, *r*^2^ of 0.1, 0.3, and 0.5, with 100, 250, and 500kb windows. LDpred provides the highest predicted *R*^2^ values, though there are sets of clumping parameters which perform similarly. In Fig. 3A, we report the *R*^2^ values for all tested clumping parameter values. Attempting to improve prediction, we imputed diploid genotypes, including missing SNPs, in our dataset (imputed diploid median coverage=0.987). Against expectation, the imputed data generally decreased accuracy compared to the unimputed data for higher coverage samples. For lower coverage samples imputation did improve *R*^2^ for the clumping approaches but not for *LDpred* (Fig. 3). One possibility is that imputation biases genetic variation to be similar to samples in the reference panel, leading to no increase in predictive power. Supporting this, if we perform MDS on the imputed genotypes, the ancestry term is no longer significant (*R*^2^ = 0.006, P=0.58), suggesting a loss of information.

**Figure 3:**
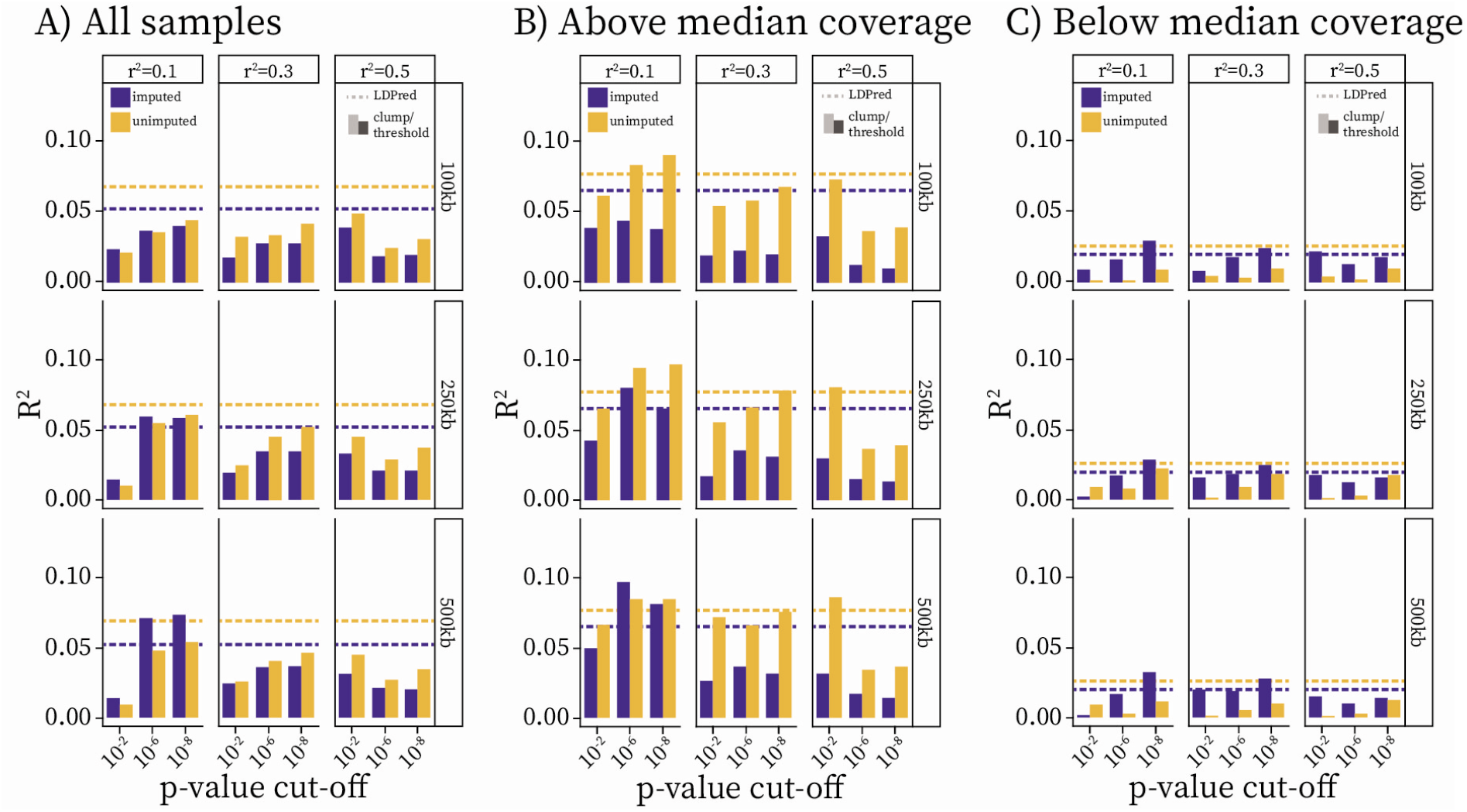
Bar plots showing the R^2^ value for each set of clumping and thresholding parameter combinations (window size, r^2^, p-value cut-off) used for PRS construction. Horizontal dashed lines show the R^2^ values from the *LDpred* PRS. Regression models include sex, date, and ancestry as covariates.

When we include geographic and climate variables in the model, the contributions of annual precipitation, mean daily temperature and latitude, are non-significant (Table 3, all < 1%). Longitude has an *R*^2^ of 3% (P=0.006; an increase of 0.07 cm/°), but its contribution to the model changes depending on which other variables are included. For instance, when ancestry is excluded from the model, longitude is no longer significant (p=0.099), suggesting a complex relationship which may obscure its effects. It is likely that this variable is a proxy for others that are not included in our models, and so this significant value is difficult to interpret. Finally, the effects of stable isotope values are not significant (Table 3), though this might be due to the relatively small sample size (n=67). Indeed, neither PRS nor ancestry is significant in this subset. Though not significant, our results suggest that increasing *δ*^15^N may be associated with increased femur length (increase of 0.4 cm /‱*δ*^15^N.

**Table 3:** Linear model results for femur length, climate and stable isotope variables

| Model | Variables | $R^2$ | permuted values | |
| --- | --- | --- | --- | --- |
| | | | $R^2$ SD | p-value |
| Femur length<br>with climate<br>and<br>geographic<br>variables<br>(n=117) | PRS | 0.0569 | 0.0066 | 0.000 |
|  | ancestry | 0.0679 | 0.0130 | 0.004 |
|  | sex | 0.1251 | 0.0079 | 0.000 |
|  | date | 0.0690 | 0.0093 | 0.000 |
|  | latitude | 0.0047 | 0.0083 | 0.381 |
|  | longitude | 0.0317 | 0.0090 | 0.019 |
|  | avgTemp | 0.0050 | 0.0065 | 0.296 |
|  | annualPrecip | 0.0074 | 0.0058 | 0.191 |
| Femur length<br>with diet<br>variables<br>(n=67) | PRS | 0.002 | 0.0117 | 0.623 |
|  | ancestry | 0.054 | 0.0245 | 0.205 |
|  | sex | 0.234 | 0.0209 | 0.000 |
|  | date | 0.032 | 0.0171 | 0.171 |
| | $\delta^{13}\text{C}$ | 0.012 | 0.0125 | 0.266 |
| | $\delta^{15}\text{N}$ | 0.030 | 0.0147 | 0.098 |

## Discussion and Conclusion

We find that that we can predict a small but statistically significant proportion of individual height variation using polygenic scores in ancient individuals. Given that our data are pseudo-haploid, meaning we observed only one chromosome of each chromosome pair, we would automatically expect our PRS predictions to perform half as well as they would on diploid data. From there, on average, about 50% of SNPS are missing in each individual, decreasing our expected predictions by another half. Therefore, we would expect to be able to predict about one quarter of the variation that can be predicted in present-day Europeans, which is roughly consistent with our findings. Surprisingly, we find that genotype imputation, at least as implemented here, does not substantially increase predictive accuracy. Further developments in imputation approaches specific to ancient DNA might provide improvements in the future. Our results show that polygenic scores cannot accurately predict individual traits, but do support their application to the quantitative study of evolutionary trends and environmental relationships on a population level (Cox et al., 2019).

There is a correlation between PRS and ancestry: genome-wide ancestry explains a similar amount of variation in height as the PRS, but when ancestry is removed, the variation accounted for by the PRS increases. There are two non-exclusive explanations for this observation. One is that genetic height varies systematically with ancestry—consistent with the observation that, on a population level, stature tracks genetically predicted height through time (Cox et al., 2019). A large portion of the predicted genetic change in stature is attributable to major admixture events, which may therefore make a substantial contribution to changes in stature over time. Differences in genetic height among populations do not necessarily indicate directional selection—substantial differences can also arise under neutrality or even stabilizing selection (Harpak & Przeworski, 2020). A second possible explanation is that ancestry is spuriously correlated with environmental variables from the GWAS population. Known as population stratification, this is a common and potentially strong source of bias in GWAS analysis, and while measures are taken to reduce its impact, there can still be evidence of residual population stratification in the GWAS results (Sohail et al., 2019; Berg et al., 2019). However, for this to affect our study it would also require a somewhat coincidental correlation between ancient and present-day stratification. With current methods and data, the signatures of residual structure and ancestry-linked variation would appear identical. However, even if the contribution of genome-wide ancestry is entirely driven by stratification, the polygenic score still explains a significant proportion of phenotypic variation beyond its interaction with ancestry.

In addition to the genetic component, dietary variables can have a substantial impact on height outcomes. Nitrogen values are mainly associated with dietary protein intake from both plant and animal sources, but are also correlated to factors such as climate (O’Brien, 2015; Scheibner, 2016), and there is an established link between protein malnutrition/undernourishment and stunting of linear growth in children (Ghosh, 2016). Given this, we would expect to see a positive trend between nitrogen values and femur length which is present, though not significant, in our data. Carbon values are more indicative of dietary plant resources, and of the terrestrial vs. marine vs. limnic provenance of food (O’Brien, 2015; Scheibner, 2016). *C*^4^ plants, such as millet, lead to significantly lowered *δ*^13^*C* values and became widespread in Western Eurasia after ca. 3000 BCE. While Palaeolithic diets were mainly terrestrial, increased variance of *δ*^13^*C* values around 10,000 BCE reflect the increased exploitation of aquatic food resources (Scheibner, 2016). Hence, our expectations for the effect of *δ*^13^*C* on height are unclear. Moreover, stable isotope values in general may co-vary to some extent with both date and climate. Another issue is that, we did not control for the bone element from which collagen samples originated. Samples might not necessarily reflect the diet of the individual during the period that is relevant for the establishment of stature. Thus, we consider the interpretation of isotope values in our study as generally representative of subsistence patterns, rather than quantitative assays of relevant diet.

Previous work found a relationship between latitude and height in Europe which we do not observe in our sample. Cox et al. (2019) suggested that the observed latitudinal trend might be genetically driven by post-Neolithic Steppe migrations; however, even if we remove the ancestry term from our model, latitude is still not sigificant. However, our sample is biased towards Northern European collections for which we found more published metrics on DNA sampled individuals. Lack of a substantial Southern European sample might explain why we do not see a relationship.

Longitude has also previously been shown to correlate with stature in the European pre-Bronze Age periods (Ruff, 2018; Cox et al., 2019), as have climate variables (Ruff, 2018). We do replicate this observation. However, this is partly driven by the relatively tall individuals from the Danube Gorges region of Southeastern Europe (12 individuals in our sample). It has been well documented that the populations of this region do not follow the same height decreases that affect the rest of the continent through history. This has often been interpreted as a genetic trait since the nutritional status and general environment in recent times has been considered less than ideal (see citations in Ruff & Holt (2018) for further discussion). In our analysis, neither genetic ancestry nor polygenic score predict this variation although there may be genetic factors that we do not capture. The trend might also be driven by environmental factors, though we do not have the data here to speculate about what factors might be involved. This motivates further study of the basis of the distinct trends in this region.

It is not currently practical to use genetic (or environmental) data to predict individual phenotypes for height or other complex traits. However, our study shows how polygenic scores can begin to separate the effects of genetics and environment on a population level. We have shown that genetic variation can independently predict stature, validating the use of polygenic scores to track evolutionary changes (Cox et al., 2019). Future work should therefore focus on compiling anthropometric, genetic, and environmental data, as our results show promise for the application of this approach on more comprehensive data. With larger samples and more detailed information about environmental covariates, more accurate quantification of the role of environment and therefore of the relative importance of genetics and environment should be possible.

## Data Availability

All ancient DNA data used in this paper are publicly available, please see their respective citations for the sources. All skeletal metrics and polygenic scores analyzed in this paper are provided in Supplementary Table 1.

## Acknowledgments

This research was supported by a Research Fellowship from the Alfred P. Sloan Foundation (FG-2018-10647), a New Investigator Research Grant from the Charles E. Kaufman Foundation (KA2018-98559) and NIGMS award number R35GM133708. Some isotope and skeletal data were collected in collaboration with Alisa Scheibner and Julia Ebert, within the Emmy-Noether-Projekt “LiVES” funded by German Research Foundation Grant RO4148/1 (PI Eva Rosenstock). The content is solely the responsibility of the authors and does not necessarily represent the official views of the National Institutes of Health or other funding sources. This research made use of the UK Biobank Resource under Application 33923.

